# Soil microbial traits shift on contrasting timescales following revegetation of former grazing lands

**DOI:** 10.64898/2026.02.02.703401

**Authors:** Timothy M. Ghaly, Vanessa J. McPherson, Vaheesan Rajabal, Mary E. Ghaly, Nicki Taws, Rachael V. Gallagher, Johannes J. Le Roux, Sasha G. Tetu

## Abstract

Revegetation is a key strategy for restoring degraded lands globally. While this process can reshape belowground microbial communities, the extent to which such changes restore soil ecosystem functions, and whether different microbial traits recover synchronously or on distinct timescales remains less clear. Understanding this is essential for evaluating restoration success, as different microbial traits underpin distinct ecosystem services, from carbon storage to plant growth promotion, and the sequence in which these are restored can inform restoration targets and provide meaningful indicators of long-term ecological recovery. To address this, we applied deep metagenomic sequencing to characterise soil microbial responses following revegetation (spanning 1-31 years prior to sampling) on grazing agricultural lands. We find that revegetation following grazing cessation drove significant and sequential shifts in dominant microbial functions and life-history strategies. Functional changes occurred in distinct phases: an early, rapid restructuring of core soil health processes, detectable as early as three years, including enrichment of nutrient retention and carbon fixation pathways, followed by a more gradual development of plant growth-promoting traits as the plant community matures. Genome-resolved analyses of nearly 500 metagenome-assembled genomes revealed a fundamental shift in dominant microbial life history strategies: from a resource-scavenging and stress tolerance profile in grazing soils to strategies that prioritise biosynthesis and growth yields in revegetated soils. These lifestyle shifts have important implications for enhancing the microbial biomass and carbon sink potential of soils. Together, these findings show that microbial functional shifts following revegetation are temporally structured – informing expectations for when key restoration targets may be achieved, and providing a practical framework for monitoring ecosystem recovery.

## Introduction

Human land use is a major driver of environmental degradation and ecosystem service loss. Agricultural grazing land, which covers more than 25% of Earth’s land surface, is subject to significant degradation through soil compaction and altered nutrient fluxes, both of which reduce plant diversity, and collectively alter ecosystem processes and functioning [1, 2]. Revegetation, the process of restoring plant species in disturbed areas, is a widely adopted strategy for the restoration of degraded lands, with agricultural land representing a key global target for this style of restoration [3]. Although the aboveground outcomes of revegetation are often visible, the impacts on belowground ecosystems, and the bi-directional feedbacks between them, are less clear.

Soil microbiomes are fundamental mediators of ecosystem processes from local to global scales, governing biogeochemical cycles, soil structure and moisture storage, and climate regulation [4–6]. Soil microbiomes drive the cycling of nutrients that underpin ecosystem health, with microbial metabolism tightly linked to aboveground biodiversity [7]. Soil microbial communities also exert strong control over carbon storage and soil-atmosphere gas fluxes through metabolic processes such as carbon and nitrogen fixation, decomposition, and the production or consumption of greenhouse gases [8, 9]. Microbial life history strategies, defined by trade-offs between growth yield, resource acquisition, and stress tolerance, also impact soil carbon dynamics [10]. Specifically, a high *growth yield* strategy promotes soil carbon storage by increasing biomass and subsequently trapping much of this carbon as stable ‘necromass’. In contrast, a focus on *resource acquisition* can drive decomposition and carbon loss by breaking down complex organic matter; while greater investment in *stress response* strategies can increase CO_2_ respiration, as energy is redirected from biomass production to maintenance and repair. In addition to impacts on biogeochemical cycling, soil microbiomes can also have significant influence on plant health and productivity, with some beneficial members having the capacity to facilitate plant nutrient uptake, abiotic stress tolerance, and disease suppression [6].

Agricultural management practices induce substantial physical, biological and chemical changes in soil that alter microbial communities [11–13]. Understanding how microbes respond to subsequent revegetation practices is needed to evaluate ecosystem-level outcomes of such efforts. Prior work has firmly established that the composition of soil microbial communities shift in response to revegetation [14–16]. Studies using 16S rRNA amplicon profiling have shown that revegetation alters the relative abundances of dominant bacterial and fungal phyla, and that these shifts broadly parallel aboveground plant community recovery [14–16]. What remains poorly resolved, however, is whether compositional change translates into functional recovery, and whether functional shifts occur synchronously across different microbial traits or on distinct timescales. This distinction matters practically, as it provides critical information regarding when soil functional capacity is restored and whether different management targets (e.g., carbon sequestration, nutrient cycling, plant growth promotion) are achieved simultaneously or sequentially.

A further unresolved question concerns the ecological strategies of microorganisms in revegetated soils. Revegetation, through the progressive restoration of plant-derived carbon inputs, root exudate diversity, and reduced disturbance, is predicted to shift the selective landscape toward yield-oriented life history strategies over time. However, whether these predicted shifts in strategies do indeed occur post-revegetation, and at what time scales remains unclear. Understanding the dominant life history strategies of revegetated soil microbiomes has direct consequences for carbon storage (e.g., via necromass production by high-yield strategists) and nutrient cycling rates, and therefore provides an ecologically meaningful indicator of restoration trajectory. Addressing these questions requires moving beyond profiling compositional change to testing mechanistic predictions about how microbial life-history strategies and ecosystem functions respond to land-use change over decadal timescales.

Here, we used deep shotgun metagenomics and genome-resolved analyses to examine how revegetation across decadal timescales reshapes the functional potential and life history strategies of soil microbial communities on former livestock grazing lands. We tested the central hypothesis that revegetation following grazing cessation alters soil microbiome functional capacity relative to actively grazed soils. We further predicted that distinct classes of functional traits and life history strategies would shift on different timescales, reflecting the sequential nature of ecosystem recovery. Our findings reveal a temporally structured microbiome response to revegetation, with implications for how belowground recovery can be monitored and interpreted in the context of broader ecosystem restoration.

## Methods

### Sample collection and processing

Sampling was conducted on six livestock grazing farms (five sheep pastures and one cattle pasture) within the Southern Tablelands and South West Slopes of New South Wales, Australia. All farms are working farms that have had plots restricted from grazing and revegetated at time scales ranging from 1 to 31 years prior to our field collections. Thus, each farm contained a paired site: an active livestock pasture and an adjacent revegetated plot. Revegetation was carried out by the not-for-profit organisation, Greening Australia. Sites were revegetated either via seeding or direct plantings, covering 8 to 20 species per site (Supplementary Table S1). Specific coordinates and establishments rates for each farm can be found in Supplementary Table S1.

Soil sampling was conducted within a 24-hour period (29-30^th^ May 2024). Within each farm, we collected triplicate soil cores from both revegetated and grazing areas. Cores were taken 10 m apart along 20 m transects. Within each farm, the paired (grazing versus revegetated) transects were located within 50 m of one another. For each sample, the top ∼1 cm of material was removed, and a soil core (5 cm wide × 10 cm deep) was collected. Cores were passed through a 2 mm sieve and homogenised in the field using ethanol-sterilised equipment. From each homogenised core, we retained four subsamples in 5 mL sterile tubes for independent DNA extractions, plus additional soil filled in a sterile 50 mL tube for physicochemical analyses. All samples were immediately placed on ice in a cooler box. Material for DNA extraction was stored at −20°C until processing, while samples for soil physicochemical measurements were kept at 4°C and processed within one week at the Environmental Analysis Laboratory (Southern Cross University, Lismore, Australia). Measured physicochemical properties included pH, electrical conductivity, available phosphorus, calcium, magnesium, potassium, effective cation exchange capacity, total carbon and total nitrogen.

Total metagenomic DNA was extracted from 0.25 g of each quadruplicate soil subsample using a previously established bead-beating protocol [17], and subsequently pooled by sample at equimolar concentrations. DNA libraries were prepared for shotgun metagenomic sequencing and run on an Illumina NovaSeq X Plus 10B (300-cycle) platform at the Ramaciotti Centre for Genomics (UNSW, Sydney, Australia), producing, on average, 137.6 million × 150-bp paired-end reads (equivalent to 20.6 gigabases) per sample. All sequence data have been made available via the European Nucleotide Archive (ENA) under project accession PRJEB97193 (ENA sample accessions: ERS26780466 – ERS26780501).

### Sequence read processing and metagenomic assembly

Paired-end metagenomic reads underwent adapter-clipping and quality-trimming using fastp v0.24.1 [18, 19], employing a 4bp sliding window, from 5’- to 3’-end, with a mean quality threshold of 20 [parameters: --cut_right --cut_right_window_size 4 --cut_right_mean_quality 20 --thread 16]. Reads were then cleaned to remove Illumina PhiX control spike-in and contaminating human sequences. Contaminant reads were identified by mapping the trimmed reads to the human telomere-to-telomere (T2T-CHM13+Y v2.0) [20, 21] and *Escherichia* phage phiX174 (NC_001422.1) genomes, using Strobealign v0.16.0 [22]. Paired-end reads where both read pairs mapped to a reference genome were identified with SAMtools [23], and removed using the *filterbyname* program from the BBTools software suite (https://sourceforge.net/projects/bbmap/). Contaminant reads accounted for 0.03% – 0.1% (median=0.06%) of reads. The final set of filtered reads were then assembled for each sample using MetaHipMer2 [24], with default settings. Each single-sample assembly was run across 40 HPC nodes on the Gadi supercomputer (National Computational Infrastructure Australia), with 1,920 CPU cores and 7.6 Tb of available memory.

### Taxonomic profiling of the metagenomes

Taxonomic profiling of the soil metagenomes was performed with SingleM v0.19.0 [25], run using the *pipe* subcommand. The SingleM *summarise* subcommand was used to obtain relative abundances for each taxonomic rank. To calculate beta-diversity between samples, we used read counts of the universal marker, S3.5 ribosomal protein S2, clustered at 96.67% using the SingleM *summarise* subcommand. The OTU counts were normalised by total-sum scaling (TSS) using the *transform_sample_counts* function from the phyloseq v1.3.2 R package [26]. Non-metric multidimensional scaling (NMDS) was used to visualise sample-level differences in community composition. First, all pairwise Bray-Curtis dissimilarities [27] were calculated from the normalised OTU table using the *distance* function from phyloseq, and ordinated with the *metaMDS* function from the vegan v2.5.7 R package [28] to generate a two-dimensional NMDS plot [parameters: distance = “bray”, k = 2, maxit = 999, trymax = 1000, autotransform = FALSE, wascores = FALSE]. Taxonomic alpha-diversity was measured using the Shannon diversity index [29], calculated from the TSS-normalised OTU table with the *diversity* function from vegan.

We used an eDNA approach to profile the plant community in each site via targeted assembly of full-length ITS sequences from the metagenomes. First, we merged paired-end reads with FLASH [30] [parameters: --max-overlap 150], which, together with the unmerged forward and reverse reads, were profiled against the EUKARYOME v1.9.4 ITS database [31] using Kraken 2 [32]. All reads classified by Kraken 2 as putative eukaryote ITS sequences were extracted using the *extract_kraken_reads.py* utility script, and co-assembled with the overlap-based assembler PenguiN Release 5-cf8933 [33] [parameters: penguin nuclassemble --min-contig-len 350 --num-iterations 50]. Full-length ITS sequences (ITS1-5.8S-ITS2) were identified and extracted using ITSx v1.1.3 [34] [parameters: -t all --only_full T]. ITS sequences were clustered into OTUs at 97% identity and 80% coverage using Vclust v1.2.3 [35], applying the Leiden clustering algorithm [36]. Plant OTUs were identified using the SINTAX function [37] of VSEARCH v2.29.1 [38] against the EUKARYOME ITS database. The presence or absence of plant ITS sequences was determined in each sample using CoverM, with a minimum read alignment identity of 97% and ITS sequence coverage of 50%. The ITS presence/absence table was collapsed at the transect-level, and putative plant community compositional differences were visualised using NMDS ordination as described above. We acknowledge limitations in using eDNA to infer plant community composition, including potential contamination from wind-blown pollen or leaves, and an inability to quantify relative abundances due to biased DNA representation from sources like fine roots. Future work will include vegetation surveys at soil sampling locations.

### Functional profiling of the metagenomes

Functional profiling of the soil metagenomes was performed using EcoFoldDB v2.0 [39], a database and pipeline for protein structure-guided annotations of ecologically relevant microbial functions. First, proteins were predicted using Pyrodigal v3.6.3 [40], a Python library binding to Prodigal [41] [parameters: -p meta -j 96]. This resulted in a total of 56,154,166 predicted proteins. We removed protein redundancy from this set, including fragments, by clustering the 56 million proteins with MMSeqs2 *Linclust* [42] at >95% identity and >99% coverage of the smaller sequence [parameters: mmseqs linclust --min-seq-id 0.95 --cov-mode 1 --cluster-mode 2 -c 0.99]. The resulting dataset was further reduced by clustering with MMSeqs2 *cluster* [43] with 90% identity and 80% bidirectional coverage thresholds [parameters: mmseqs cluster --min-seq-id 0.9 --cov-mode 0 -c 0.8], generating a non-redundant set of 29,725,552 proteins. These protein sequences were functionally annotated using the *EcoFoldDB-annotate* v2.0 pipeline with default settings, which leverages the ProstT5 protein language model [44] and Foldseek [45] to allow rapid protein structure-based functional annotations against EcoFoldDB directly from sequence data.

### Genome binning and analysis of metagenome-assembled genomes

Assembled contigs were binned into clusters of metagenome-assembled genomes (MAGs) using GenomeFace [46]. Prior to binning, contig coverage calculation was performed using CoverM v0.7.0 [47], which mapped the filtered reads from each sample to the complete set of pooled contigs using Strobealign [parameters: contig -p strobealign --strobealign-use-index --min-read-percent-identity 97 -m metabat]. The contig coverage information was provided to GenomeFace for binning in ‘single sample assembly’ mode, and with a minimum contig length of 1,000 bp [parameters: -s -m 1000]. Genome quality of the resulting MAGs was assessed using CheckM2 [48]. Only high-purity MAGs, with CheckM2-estimated contamination <5% and completeness > 50%, were retained for downstream analysis. Filtered MAGs were then dereplicated at 99% average nucleotide identity (ANI) using Galah [49], as part of the CoverM package. Galah was run using skani [50] for ANI calculations, and the Checkm2 output for genome quality information [parameters: coverm cluster --ani 99 --checkm2-quality-report --precluster-method skani --cluster-method skani].

MAGs were taxonomically classified using GTDB-Tk v2.4.1 [51, 52] against GTDB release 226 [53–56] [parameters: classify_wf --skip_ani_screen]. Domain-specific phylogenies were inferred from the concatenated protein alignments produced by GTDB-Tk, based on the *bac120* [57] and *ar53* [58] marker sets. Approximate maximum-likelihood phylogenetic trees were constructed from the alignments using FastTree2 v2.1.11 [59], applying the WAG amino acid substitution model [60], and with branch lengths rescaled to optimise the Gamma20 likelihood [parameters: -gamma -wag]. The trees were rooted with minimal ancestor deviation (MAD) [61] using MADroot [62] (https://github.com/davidjamesbryant/MADroot), and visualised using the *ggtree* [63] and *ggtreeExtra* [64] R packages.

Metabolic pathways encoded by the MAGs were identified using MetaPathPredict [65], which takes KofamScan annotations [66], and applies machine learning models to predict the presence or absence of KEGG modules [67] within incomplete genomes. For trophic-lifestyle inference, each MAG was evaluated for carbon-fixation pathways (based on MetaPathPredict and EcoFoldDB annotations), and for substrate profiles of carbohydrate-active enzyme (CAZyme) gene clusters (CGCs) using the run_dbcan v5.1.2 [68] *easy_substrate* pipeline. MAGs encoding carbon-fixation pathways but lacking detectable CGCs were classified as autotrophs; MAGs with CGCs but no carbon-fixation pathways were classified as heterotrophs; and MAGs with both features were classified as mixotrophs.

Genome-based maximum growth rates were predicted for each MAG using Phydon [69], which incorporates codon usage biases and phylogenetic signal for robust predictions of minimum doubling times. Genome annotation data used by Phydon was generated using Prokka v1.14.6 [70], while the evolutionary relationship data were based on the phylogenetic placements of each MAG in the GTDB bacterial and archaeal trees, produced by GTDB-Tk.

### Statistical analyses

All statistical analyses were conducted in R [71]. To control the false discovery rate (FDR) arising from multiple hypothesis testing, all *p*-values were adjusted using the Benjamini-Hochberg method [72] where applicable.

We used the MaAsLin 2 R package [73] to identify OTUs differentially abundant between grazing and revegetated soils. Linear models were fit with land-use (grazing vs revegetated) as a fixed effect, and farm as a random effect. OTU counts were normalised by total-sum scaling (TSS) and log-transformed. OTUs present in fewer than 10% of samples were excluded.

MaAslin 2 was also used to identify MAGs enriched in either grazing or revegetated soils. MAG per-base abundances were estimated with CoverM in genome mode (parameters: genome -p strobealign -m mean), and then scaled for sequencing depth by the average number of read pairs across all samples. These normalised MAG abundances were analysed with MaAsLin2 using the same model structure as the OTU analysis, but with MaAsLin2 count normalisation disabled.

We used linear modelling to identify microbial functions enriched in revegetated versus grazing soils. Gene relative abundances were taken from the CoverM-calculated coverage of their contigs (i.e., per-base contig abundances scaled for sequencing depth by the average read-pair count), and summed for each EcoFoldDB functional category. We fit the model: *Abundance ∼ Farm + Land-use + Land-use × Time*, for each function category where, *Abundance* = summed normalised abundance of that function; *Farm* = blocking covariate accounting for paired sampling design (i.e., grazing and revegetated transects from the same farm); *Land-use* = grazing versus revegetated; and *Land-use × Time* = interaction term of years since revegetation (mean-centred).

Alpha-diversity (Shannon index) differences were tested with Wilcoxon rank-sum tests [74]. Differences in beta-diversity (Bray-Curtis dissimilarities) were tested with PERMANOVAs using the *adonis2* function from the vegan R package. We applied environmental vector fitting to examine any associations between measured soil abiotic variables and microbial community composition using the *envfit* function from vegan. Fisher’s exact tests [75] were used to identify KEGG modules and trophic-lifestyle categories over-represented in revegetated-versus grazing-enriched MAGs. Linear models were used to test for correlations between MaAsLin2 MAG enrichment coefficients and Phydon-predicted maximum growth rates.

## Results and Discussion

### Shifts in the compositional structure of soil microbiomes

We used deep shotgun metagenomic sequencing to assess soil microbiome responses to revegetation following the cessation of livestock grazing. Average soil microbial biodiversity, assessed using the Shannon diversity index (alpha-diversity), did not differ significantly between grazed and revegetated sites, although, the variation in diversity was greater in revegetated soils (Fig. 1b; Supplementary Fig. S1). In contrast, beta-diversity was found to be significantly different (Fig. 1b, c), indicating that revegetation following the cessation of grazing significantly altered microbial community composition relative to grazed soils. Notably, this restructuring occurred independently of major abiotic soil properties, as no measured physicochemical parameter was significantly correlated with community composition via environmental fitting (FDR-adjusted *p*-values > 0.05), although a trend towards lower pH was observed in revegetated soils (Supplementary Fig. S2). We found that soils from sheep-grazed pastures shifted as early as 3 years post-revegetation, forming a distinct cluster across the 3-to-31-year range (PERMANOVA: sheep-grazed versus revegetated samples, *p* = 0.001). In contrast, the only cattle-grazed pasture (F3) exhibited no compositional shift, even after 11 years since revegetation (Fig. 1c; PERMANOVA: cattle-grazed versus revegetated samples, *p* = 0.1). This differential response might indicate that the restoration of grazing soil microbiomes is livestock-specific; an effect possibly stemming from cattle causing a higher degree of initial soil degradation than sheep, though further investigation is required to confirm this. In contrast to these observed soil microbiome shifts, we found a consistent convergence in putative plant community composition (inferred from eDNA) regardless of previous land use, with all revegetated sites >3 years old clustering together and differing significantly from grazed pastures (PERMANOVA, *p* = 0.008; Supplementary Fig. S3).

**Fig. 1.**
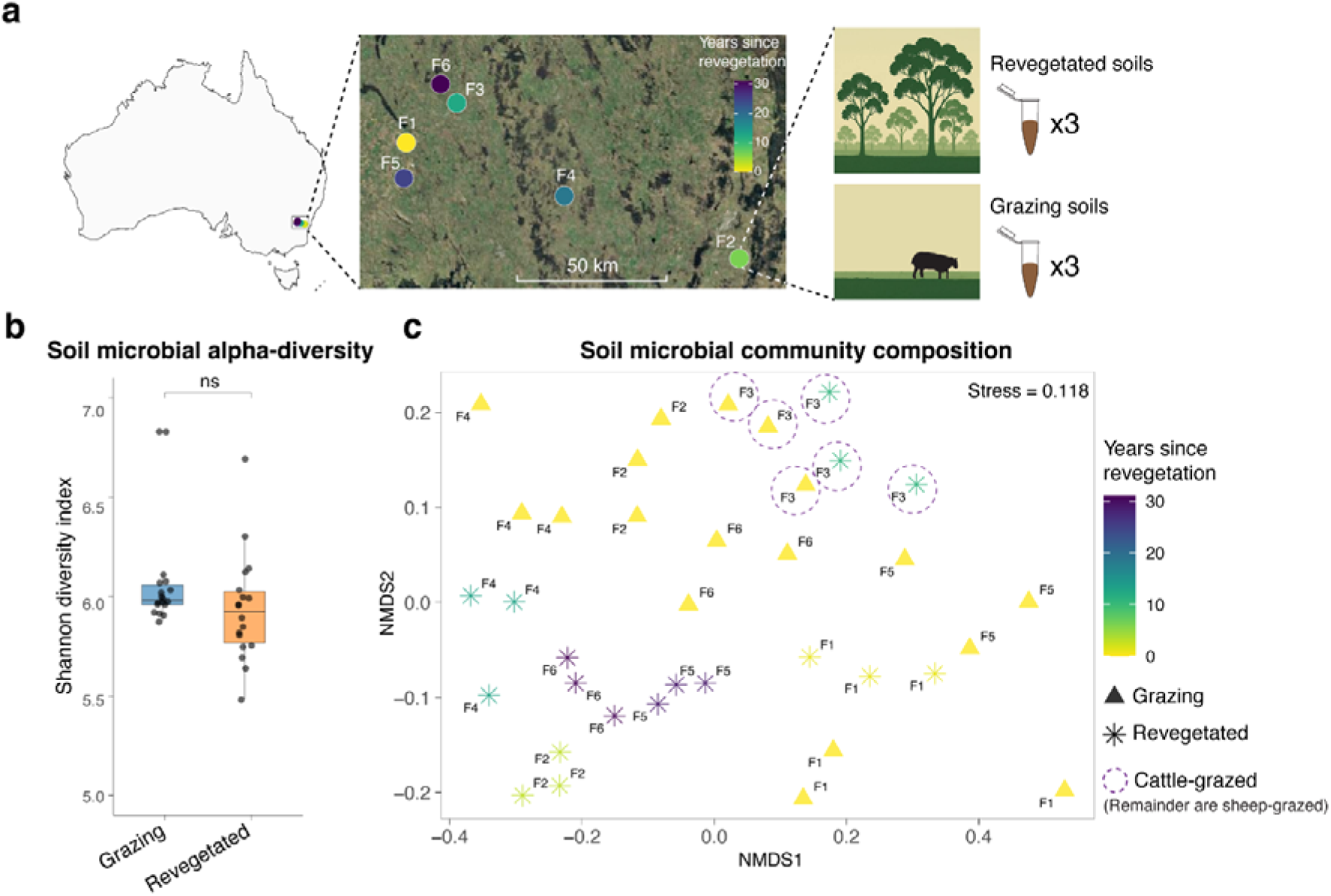
The impacts of revegetation following grazing cessation on soil microbial community diversity. **a**, Soil was sampled from six active livestock farms in NSW, Australia. Revegetated areas had been restored 1-31 years prior. At each farm, triplicate soil cores were sampled from both grazing and adjacent revegetated areas. **b**, Soil microbial alpha-diversity (Shannon index) comparisons between grazing (blue) and revegetated soils (orange). See Supplementary Fig. S1 for farm-wise comparisons. **c**, Non-metric multidimensional scaling (NMDS) visualising sample-level differences in soil community composition (i.e., beta-diversity). Community dissimilarities are derived from pairwise Bray-Curtis distances. Point shapes indicate land-use type: grazing (triangles) versus revegetated (stars). Points are coloured by years since revegetation. Text labels indicate the farm from which the samples derived, as shown in panel **a**. Note, F3 represents the cattle-grazed farm, while all others are sheep-grazed.

Revegetated soil microbiomes were characterised by a consistent increase in the relative abundance of Actinomycetota (Fig. 2a), the dominant phylum across all samples. This shift was driven by the enrichment of specific Actinomycetota taxa at the OTU level (Fig. 2b), particularly from the family Streptosporangiaceae, as well as members of the Solirubrobacteraceae, Pseudonocardiaceae and Nocardioidaceae families. Other significantly enriched OTUs belonged to the genus *Bradyrhizobium* (class Alphaproteobacteria) (Fig. 2b), the predominant rhizobial genus that forms symbioses with many Australian native legumes [76].

**Fig. 2.**
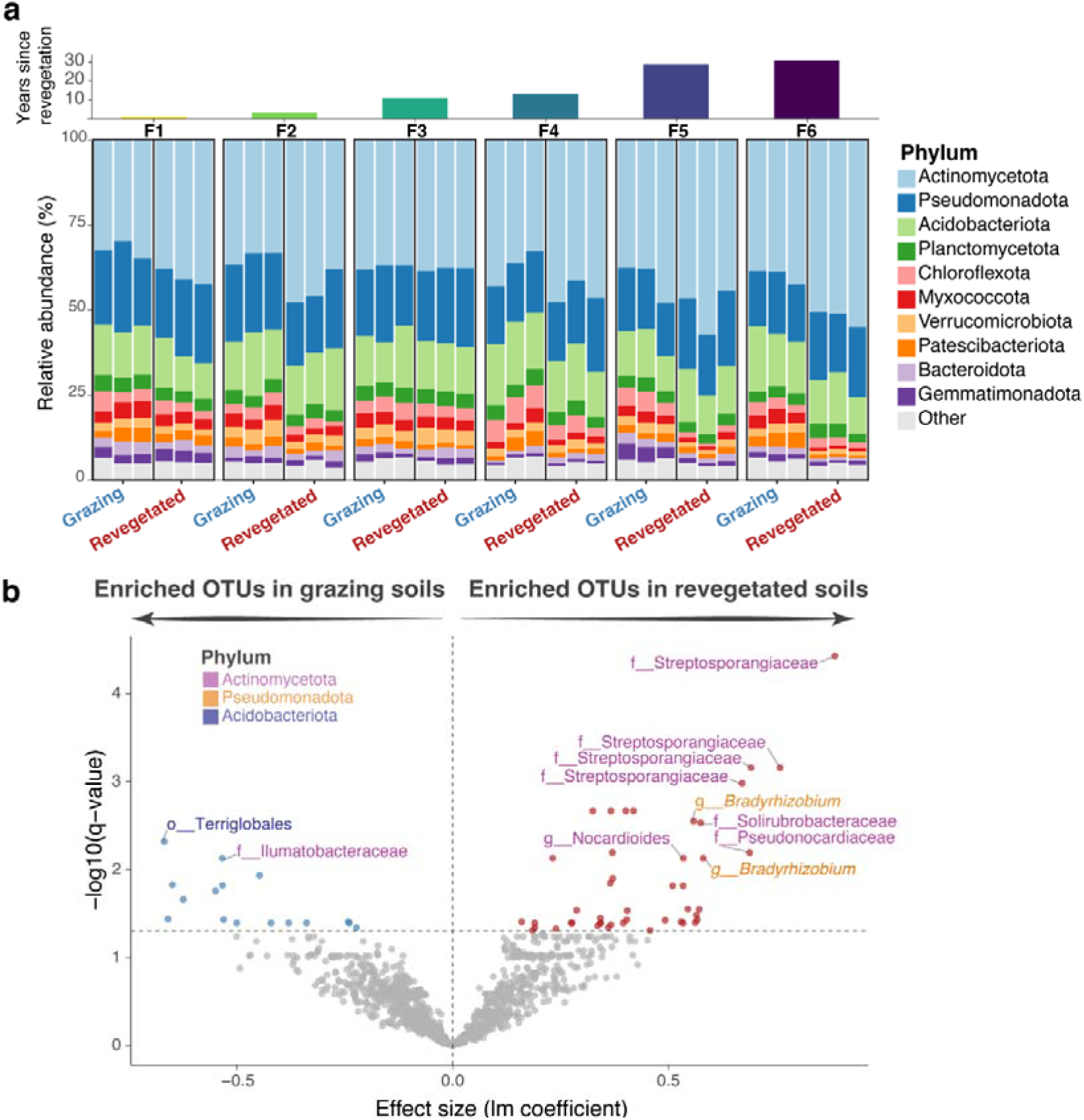
Taxonomic shifts in soil microbiomes following grazing cessation and revegetation. **a**, Stacked bar chart shows phylum-level composition of soil microbial communities from each sample. The corresponding farm and years since revegetation are shown above. **b**, Microbial OTUs significantly enriched in grazing versus revegetated soils. The most significant OTUs, with an enrichment coefficient > 0.5 and *q*-value (FDR-adjusted *p*-value) < 0.01, are labelled with their lowest taxonomic classification, and coloured by phylum.

### Shifts in the functional potential of soil microbiomes

Cessation of grazing and revegetation drove significant shifts in the functional potential of the soil microbial communities (Fig. 3). Using protein structure-based functional profiling with EcoFoldDB [39], we identified genes linked to key ecological traits. Soils revegetated after cessation of grazing were enriched relative to grazed soils in functions associated with a suite of diverse biogeochemical processes, including carbon fixation, nutrient mineralisation and retention (N, P and S) and plant-microbe interactions (Fig. 3).

**Fig. 3.**
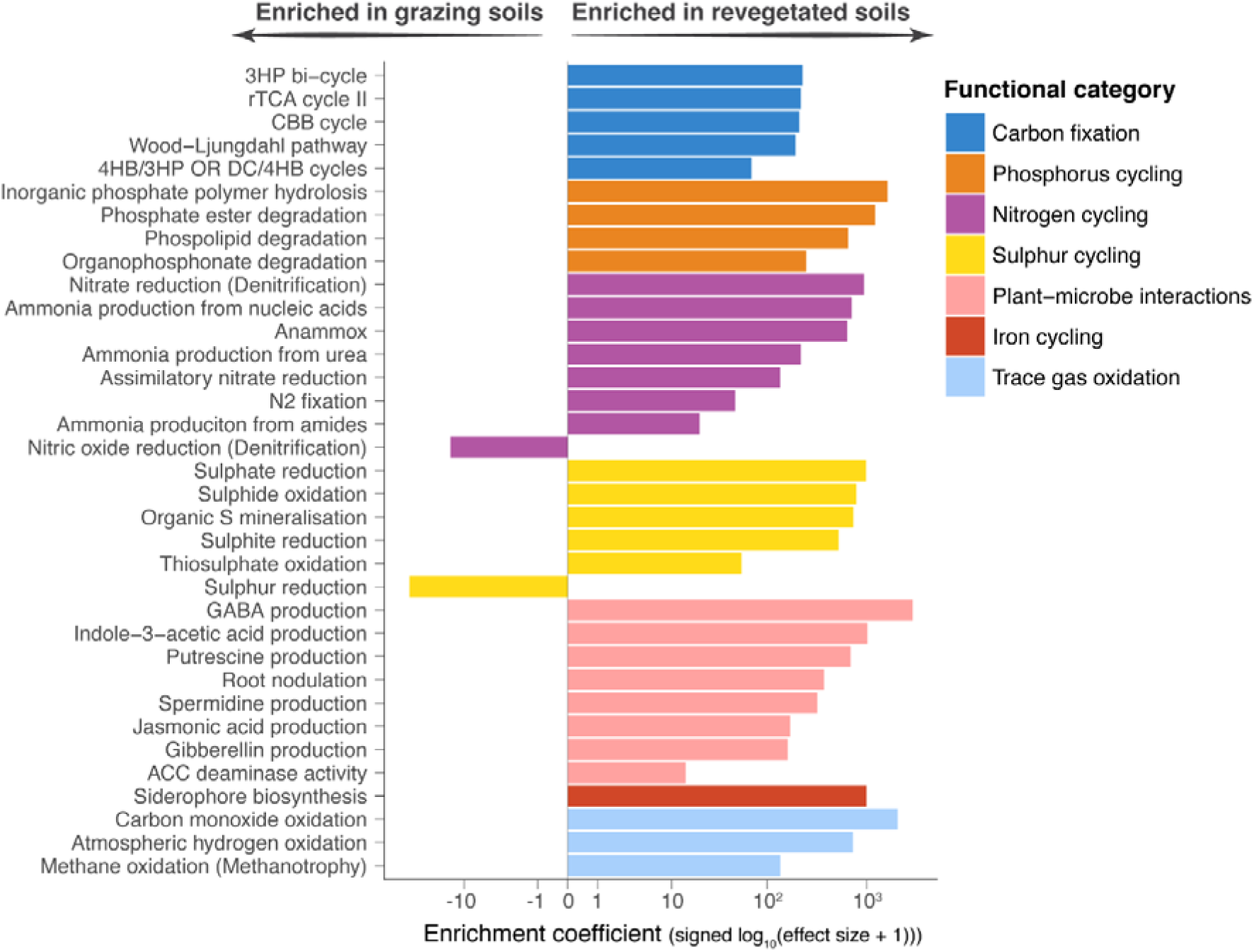
Functional shifts in soil microbiomes following revegetation. Community functions that are significantly enriched (FDR-adjusted *p*-values < 0.05) in grazing versus revegetated soils. Bars are coloured according to their broader functional category.

All of the core carbon fixation pathways present in prokaryotes [77] were significantly enriched in the soil microbial communities of revegetated sites, indicating a heightened potential for soil carbon sequestration. As revegetation is strongly associated with increased soil organic carbon storage [78, 79], our findings suggest that the enrichment of diverse carbon-fixing taxa may play a key role in this re-carbonisation of soils. In addition, inorganic phosphorus solubilisation and organic phosphorus mineralisation were enriched in revegetated soils, suggesting a community with a greater potential for increasing plant-available phosphorus. We also observed nitrogen cycling pathways enriched in the revegetated communities, involving nitrogen acquisition and recycling. These included assimilatory nitrate reduction (incorporating nitrogen into biomass), nitrogen fixation (introducing new nitrogen), and the recycling of organic nitrogen compounds (e.g., nucleic acids, amides, urea) into plant-available ammonia. Revegetated soils were also enriched in pathways representing complete sulphur cycling, including the mineralisation of organic sulphur, reduction of sulphate, and the re-oxidation back to sulphate. This sulphur cycling is likely to sustain plant-available sulphur and prevent the accumulation of toxic hydrogen sulphide.

In contrast, soils which continue to be grazed were significantly enriched in nitric oxide reduction, relative to those revegetated, a central step in denitrification that promotes the direct loss of nitrogen from soil systems through the emission of nitrous oxide (N □ O). This also highlights a potential climate change related consequence associated with grazed soils since N □ O is a potent greenhouse gas with a warming potential approximately 300 times that of CO □ [80]. Additionally, grazing soils were enriched in sulphur reduction to hydrogen sulphide (without the corresponding enrichment of the complementary oxidative pathways required for a complete sulphur cycle). This has the potential to increase hydrogen sulphide accumulation, which can inhibit other microbial processes as well as plant growth.

Soil microbiomes within revegetated areas were also significantly enriched in genes involved in beneficial plant-microbe interactions (Fig. 3). Key enriched functions included ACC deaminase activity, which modulates plant ethylene levels to alleviate abiotic stress [81]; the synthesis of phytohormones such as indole-3-acetic acid (auxin), jasmonic acid, and gibberellins to directly stimulate plant growth and defence [82–84]; and the production of polyamines (e.g., putrescine and spermidine) and gamma-aminobutyric acid (GABA), which contribute to stress resilience and root development [85, 86]. Furthermore, the enrichment of root nodulation genes highlights the shift in microbial community to support mutualistic symbioses between rhizobia and native legumes. Collectively, these findings indicate that the cessation of grazing together with revegetation drive functional shifts in soil microbiomes towards processes important for soil, plant, and ecosystem health.

### Temporal dynamics of shifts in microbial function

A key finding of our study is the distinct chronology of microbial functional shifts following the cessation of grazing and revegetation. The time-series analysis revealed that soil microbiome responses to grazing cessation and revegetation occurs in two broad temporal phases. We found that many of the functions observed to be enriched in the revegetated soils had no correlation with time, indicating that fundamental shifts in biogeochemical cycling occur rapidly (within ∼3 years) and then stabilise over longer time scales. This suggests a rapid initial restructuring of core ecosystem processes, such as carbon fixation and nutrient mineralisation, establishing a new, stable functional baseline soon after land use changes.

However, a second set of functional shifts occurred more gradually following grazing cessation and revegetation (Fig. 4). Among the functions that were significantly correlated with age in these sites, those facilitating plant-microbe interactions represented the most abundant category. This pattern suggests that as the plant community grows and matures over time, selection for more specialised plant-microbe interactions increases along with it. Consequently, the development of a native plant-adapted microbiome appears to be an ongoing process that continues to develop over longer (decadal) periods following revegetation. This decadal-scale maturation may reflect a gradual fine-tuning of the soil microbiome, aligning with the establishing plant community, however, specific tests of this are needed. Beyond plant-associated functions, several other metabolic processes showed progressive enrichment with time since grazing cessation and revegetation, including urease activity, phospholipid degradation, thiosulphate oxidation, and methane and carbon monoxide oxidation, suggesting community adaptation to more diverse metabolic niches as revegetated sites develop.

**Fig. 4.**
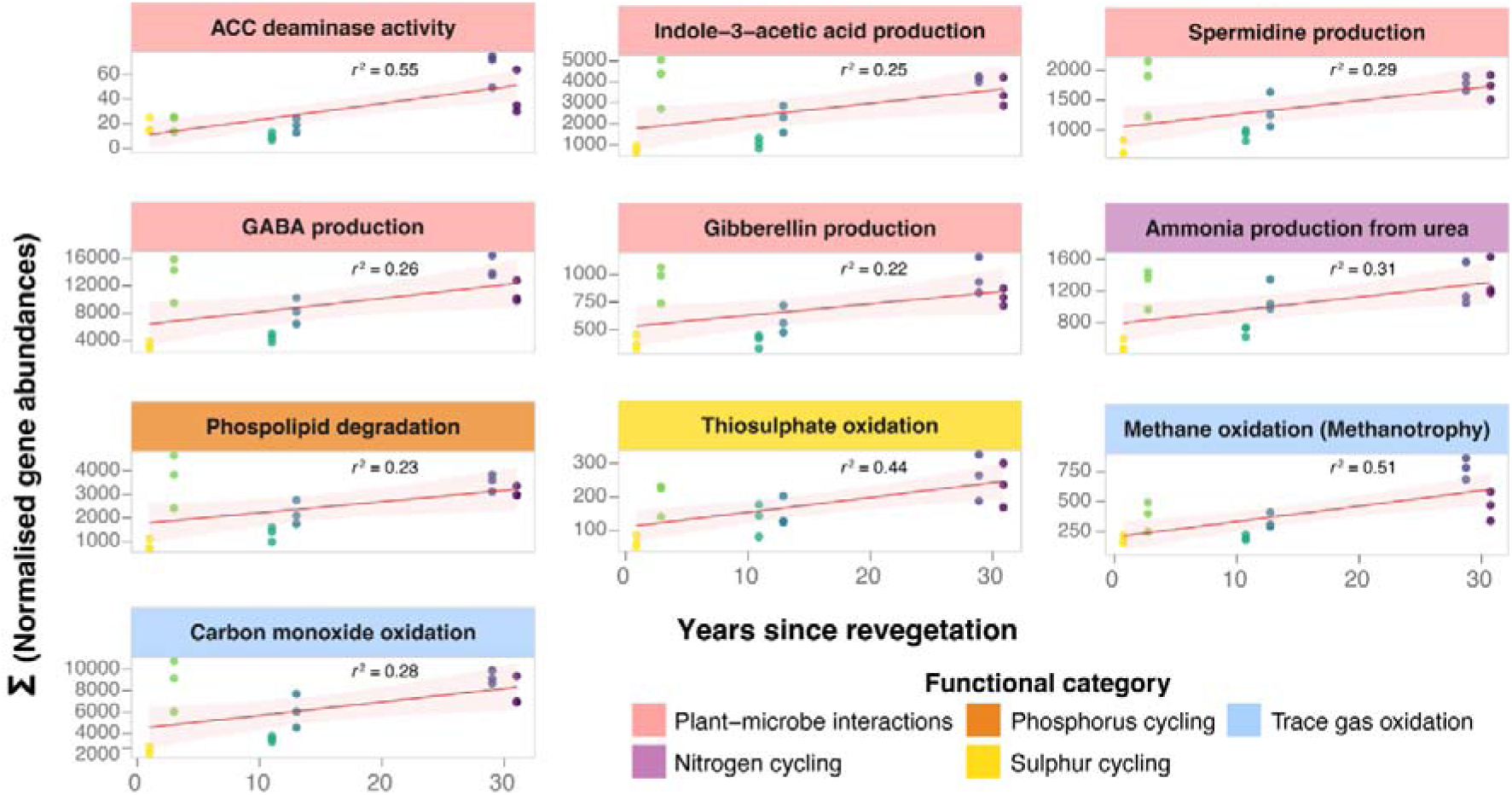
Functional shifts correlated with time since revegetation. Individual panels show community functions (summed normalised gene abundances) that are significantly correlated (FDR-adjusted *p*-values < 0.05) with years since revegetation and cessation of grazing. Panels are coloured according to their broader functional category.

Together, these findings suggest that the restoration of soil microbiome function after grazing cessation and revegetation follows a clear temporal sequence: an initial, rapid transformation of core biogeochemical cycles, followed by a prolonged, gradual specialisation towards more complex plant-microbe interactions and niche adaptation.

### Shifts in microbial life history strategies

We next performed genome-resolved analyses to assess the impact of grazing cessation along with revegetation on genomic traits linked to microbial growth and life history strategies. From the assemblies, we reconstructed 442 bacterial and 12 archaeal MAGs of high purity (>50% completeness and >95% purity), primarily representing the phyla Actinomycetota, Acidobacteriota, and Pseudomonadota. We observed a phylogenetic signal in the patterns of genome enrichment, indicating that responses to grazing cessation and revegetation is often lineage-specific (Fig. 5). This suggests that evolutionary history influences microbial enrichment, likely stemming from certain clades sharing conserved functional or physiological traits that drive their success in either grazing or revegetated soils.

**Fig. 5.**
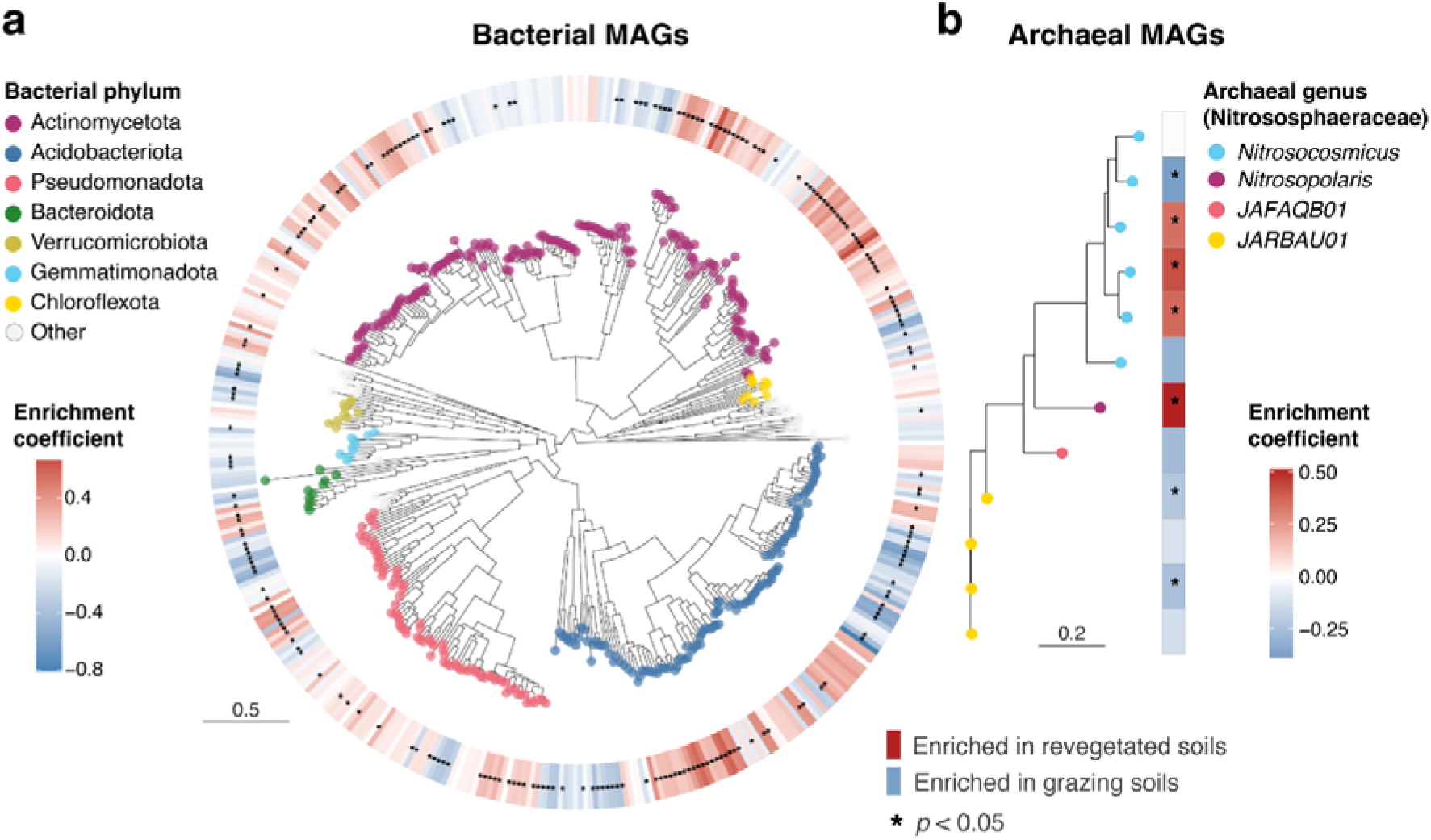
Enrichment of metagenome-assembled genomes (MAGs) in grazing and revegetated soils. Phylogeny of **a**, 442 bacterial and **b**, 12 archaeal high-purity (>95% purity, >50% completeness) MAGs. Tree tip points are coloured by GTDB phylum. Scale bars indicate amino acid substitutions per site. Colour scale bars indicate enrichment in grazing (blue) versus revegetated (red) soils, and asterisks indicate statistical significance (FDR-adjusted *p*-values < 0.05).

Grazing cessation coupled with revegetation significantly altered patterns of microbial life history traits. Growth rate predictions, based on codon usage bias and phylogenetic signal (which serve as robust predictors for microbial growth potential [69, 87]), revealed a significant enrichment of faster-growing taxa in revegetated soils (Fig. 6a). This aligns with previous observations of copiotrophic (faster-growing) taxa enriched in soils following revegetation [88, 89], potentially driven by increased inputs of plant-derived organic matter. Our results, consistent with these studies, suggest that the cessation of grazing and revegetation select for microbial communities with greater investment in growth capacity. We also found that revegetated soils favoured a higher abundance of carbon-fixing taxa (Fig. 6b), key drivers of soil carbon sequestration, as well as genomes enriched in diverse biosynthetic pathways (Fig. 6c). These included the *de novo* synthesis of amino acids (e.g., tryptophan, lysine, valine, leucine), nucleotides (deoxyribonucleotides, pyrimidines), and essential cofactors (riboflavin, NAD, Coenzyme A, betaine). Collectively, this profile indicates communities geared toward enhanced synthesis of complex macromolecules, growth, and high biomass yield. This is supported by previous studies involving empirical measurements of soils, which show strong associations between revegetation and increased soil microbial biomass and soil organic carbon [78, 79, 88–92]. Our findings suggest that these increases could be driven by shifts in life history strategies that prioritise carbon fixation, biosynthesis and growth yields. This could have significant implications for enhancing the carbon sink potential of soils, warranting more targeted investigations into the effects of land use and revegetation on microbiome-mediated carbon dynamics.

**Fig. 6.**
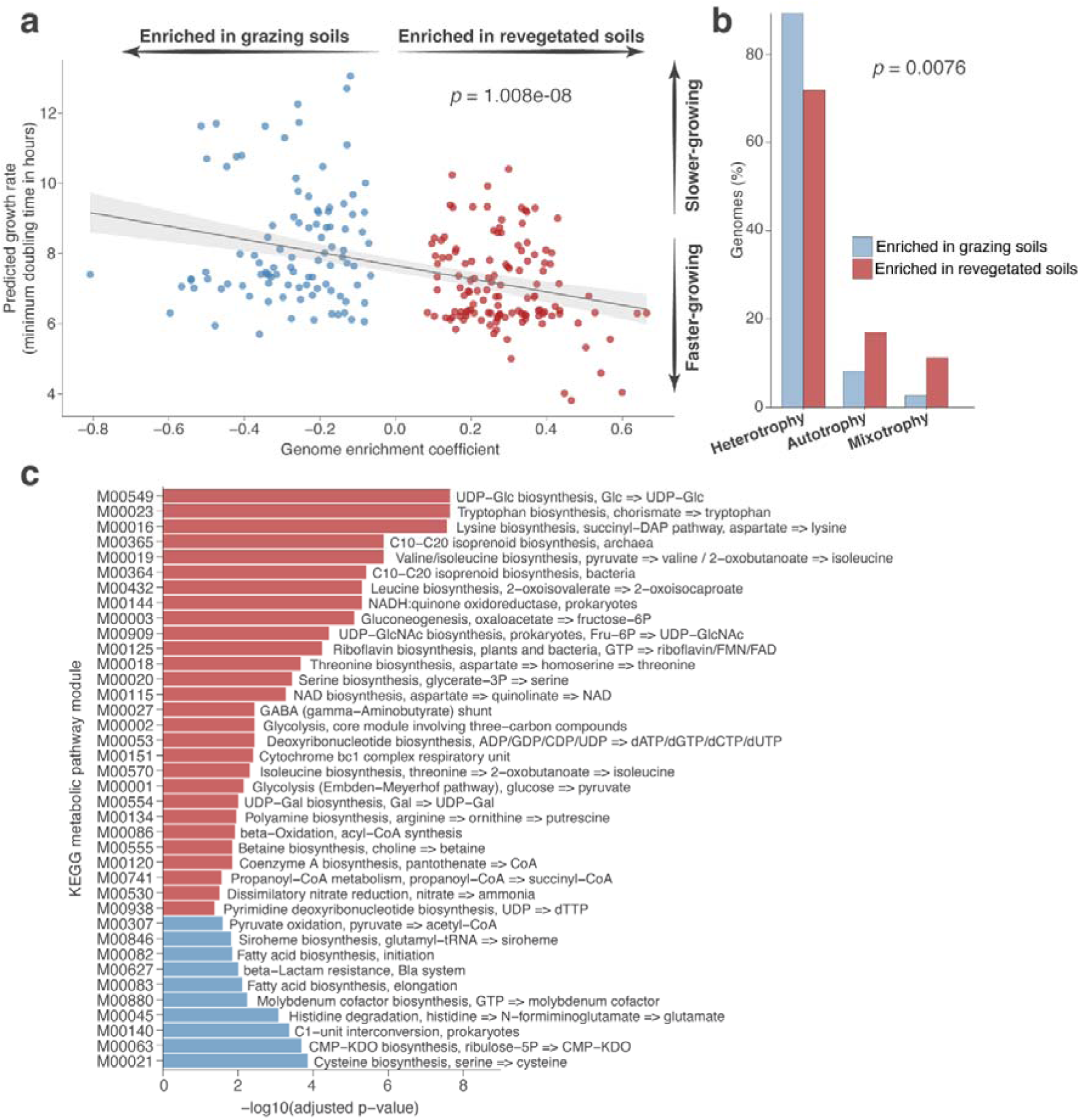
Genome-resolved enrichment of microbial life history traits. **a**, Relationship between enrichment coefficients for the metagenome-assembled genomes (MAGs) (positive = enriched in revegetated soil; negative = enriched in grazing soils) and their predicted maximum growth rates. The black line shows the linear regression with 95% confidence interval (shaded). The *p*-value for the association is displayed on the panel. **b**, Proportion of MAGs enriched in revegetated (red) or grazing (blue) samples classified into trophic categories. Bars show the percentage of MAGs in each category. The *p*-value for the difference between groups is indicated on the panel. **c**, Enrichment of metabolic pathways (KEGG modules) significantly enriched (FDR-adjusted *p*-value < 0.05) in revegetated-associated (red) versus grazing-associated (blue) MAGs. KEGG module annotations are displayed on the plot.

Conversely, grazing soils were enriched in slower-growing taxa (Fig. 6a) with a greater investment in catabolic and stress resistance mechanisms (Fig. 6b,c), suggesting a trade-off in life history strategies, favouring resource scavenging and stress tolerance, over biomass production. Grazing-enriched metabolic pathways included energy-harvesting (e.g., pyruvate oxidation, histidine degradation) and processing one-carbon compounds (C1-unit interconversion). Metabolic pathways often associated with stress response strategies were also enriched. These included beta-lactam resistance (a maker for antimicrobial and general cell envelope stress response [93]), fatty acid biosynthesis (important for maintaining cellular integrity and cell membrane repair during stress), and cysteine biosynthesis (a key pathway for oxidative stress resistance through its role in antioxidant production [94, 95]). Such investments into cellular maintenance and stress tolerance mechanisms, generally reduce microbial carbon use efficiency, diverting carbon away from growth and limiting the formation of microbial biomass and soil organic carbon [10].

## Conclusion

Here, we demonstrate that the cessation of grazing, together with revegetation, drives significant shifts in the structure, function and prevailing life history strategies of soil microbial communities. Microbiome responses were consistent across all sites where revegetation and the cessation of grazing occurred at least three years prior, with one notable exception. We note that this outlier site, which showed no significant response after 11 years, was the only previously cattle-grazed pasture, with all others being previously sheep-grazed. We hypothesise that degradation associated with cattle grazing may be more impactful on soil microbiomes, imposing a longer-lasting negative legacy that impedes microbial recovery, though further research is needed to confirm this. Despite this exception, soil microbiomes from revegetation sites were consistently associated with clear shifts relative to the sheep-grazed soil microbiomes.

This grazing-to-revegetation transition was also marked by fundamental functional shifts in the soil microbiome. In grazed soils, the microbial community was enriched in pathways involved in nitrogen loss, production of the potent greenhouse gas, nitrous oxide, and sulphur reduction, which can lead to the accumulation of toxic hydrogen sulphide. These enriched functions were accompanied in grazed soils by prevailing life history strategies focused on catabolic resource scavenging and stress-response. In contrast, revegetated soils were enriched in communities geared toward higher growth yield strategies and metabolic processes that underpin biosynthesis, carbon sequestration, nutrient retention, and plant mutualisms.

Crucially, while shifts in key biogeochemical functions linked to soil health occurred rapidly and stabilised following the cessation of grazing and revegetation, plant growth-promoting traits increased more gradually, likely in response to the progressive maturity of the plant community over time. This indicates a multi-stage microbial successional process: an early restructuring of core soil health processes, followed by a progressive development of plant-microbe interactions as the plant community matures. This multi-stage trajectory not only shows that different microbial functions shift on distinct timescales, but also carries practical implications: it informs expectations for when different management targets (such as carbon sequestration, nutrient cycling, and plant growth promotion) may be achieved following restoration efforts. This can provide meaningful indicators of restoration trajectory, informing how ecosystem restoration outcomes are monitored and interpreted.

## Availability of data and material

Raw shotgun metagenomic sequence data generated in this study have been made available via the European Nucleotide Archive (ENA), under project accession PRJEB97193 (ENA sample accessions ERS26780466 – ERS26780501).

## Authors’ contributions

T.M.G. conceived and designed the study, secured funding, contributed to fieldwork, conducted data analysis, and wrote the original manuscript draft. V.M.J. contributed to fieldwork and performed laboratory analyses. V.R. conducted laboratory work. N.T. identified sampling sites. R.V.G., J.L.R., M.E.G., and S.G.T. contributed to project design and management. All authors reviewed and edited the final manuscript.

## Acknowledgements

TMG and MEG would like to thank Saoirse and Maebh Ghaly for their loving support. The authors would also like to thank all landholders for permitting access to collection sites. We acknowledge the Traditional Custodians of the lands on which soils were sampled, the Ngunnawal and Gundungurra peoples, and pay our respects to their Elders past, present and emerging. We recognise their continuing connection to Country and the role of Indigenous land stewardship in maintaining healthy soils and ecosystems for millennia.

