## Supplementary Figs. S1-S3 for "Soil microbial traits shift on contrasting timescales following revegetation of former grazing lands"

**The PDF file includes:**

Supplementary Figs. S1-S3

**Other Supplementary Data for this manuscript include the following:**

Supplementary Table S1

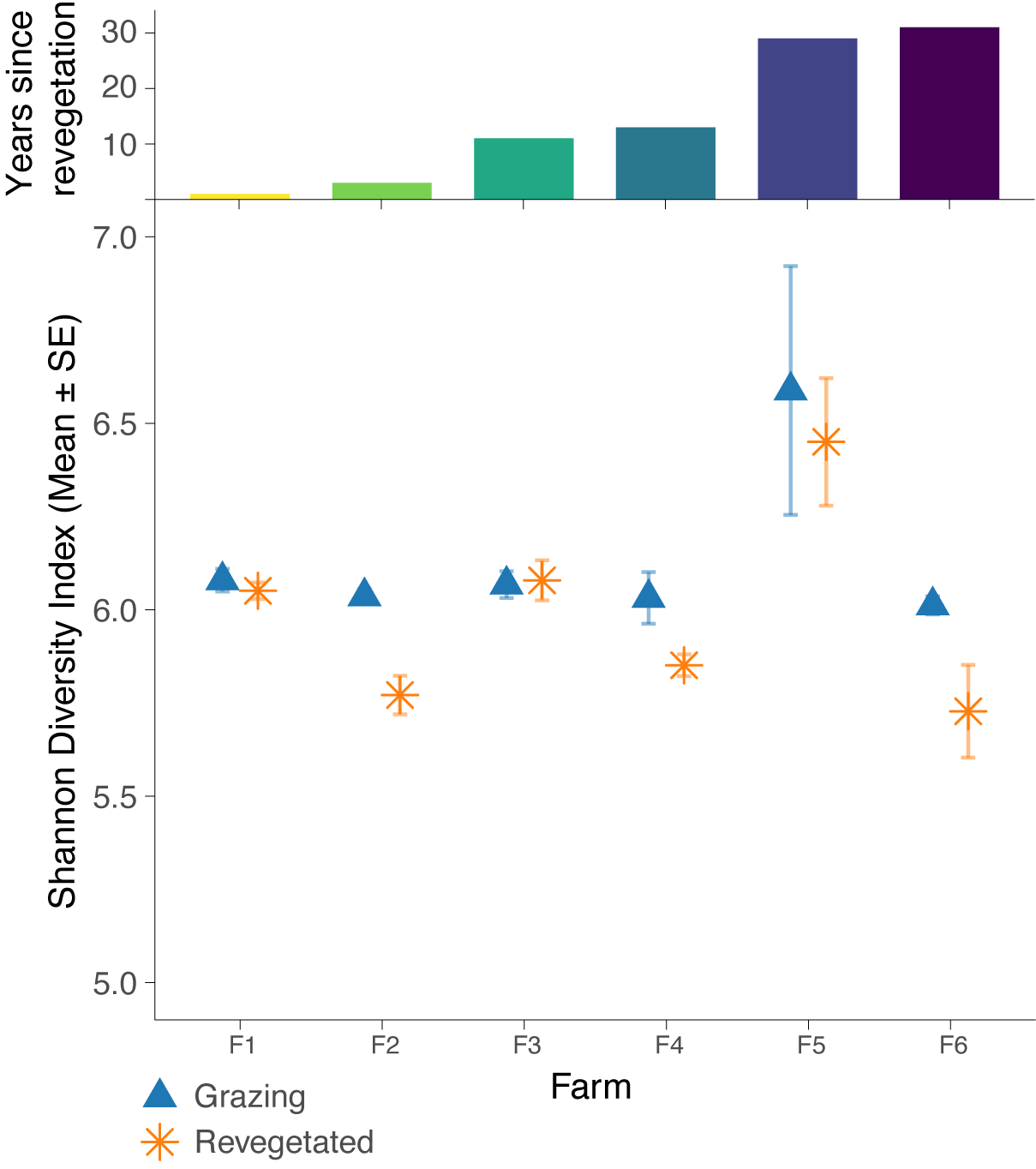

**Fig. S1. The impacts of revegetation following grazing cessation on soil microbial alpha-diversity.** Dot plots show the Shannon Diversity Index (mean ± 1 SE) for grazing (blue triangles) and revegetated (orange asterisks) soils from each farm (F1 to F6). The corresponding years since revegetation for each farm are shown as a bar chart above.

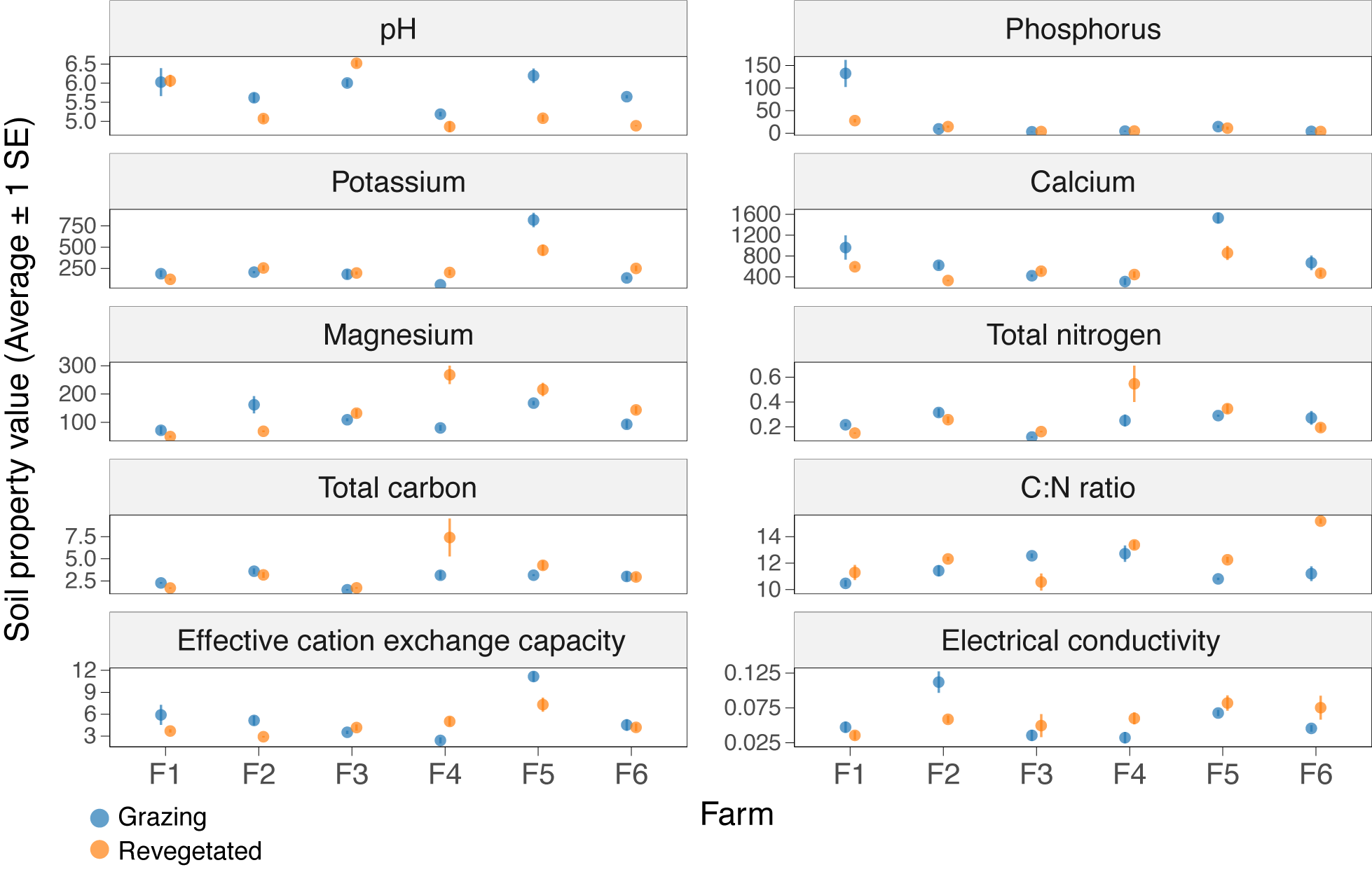

**Fig. S2. Abiotic properties of sampled soils.** Dot plots show the average soil property value (± 1 SE) for grazing (blue) and revegetated (orange) samples from each farm (F1 to F6). No significant difference was observed for any measured variable, although a trend towards lower pH was observed in revegetated soils.

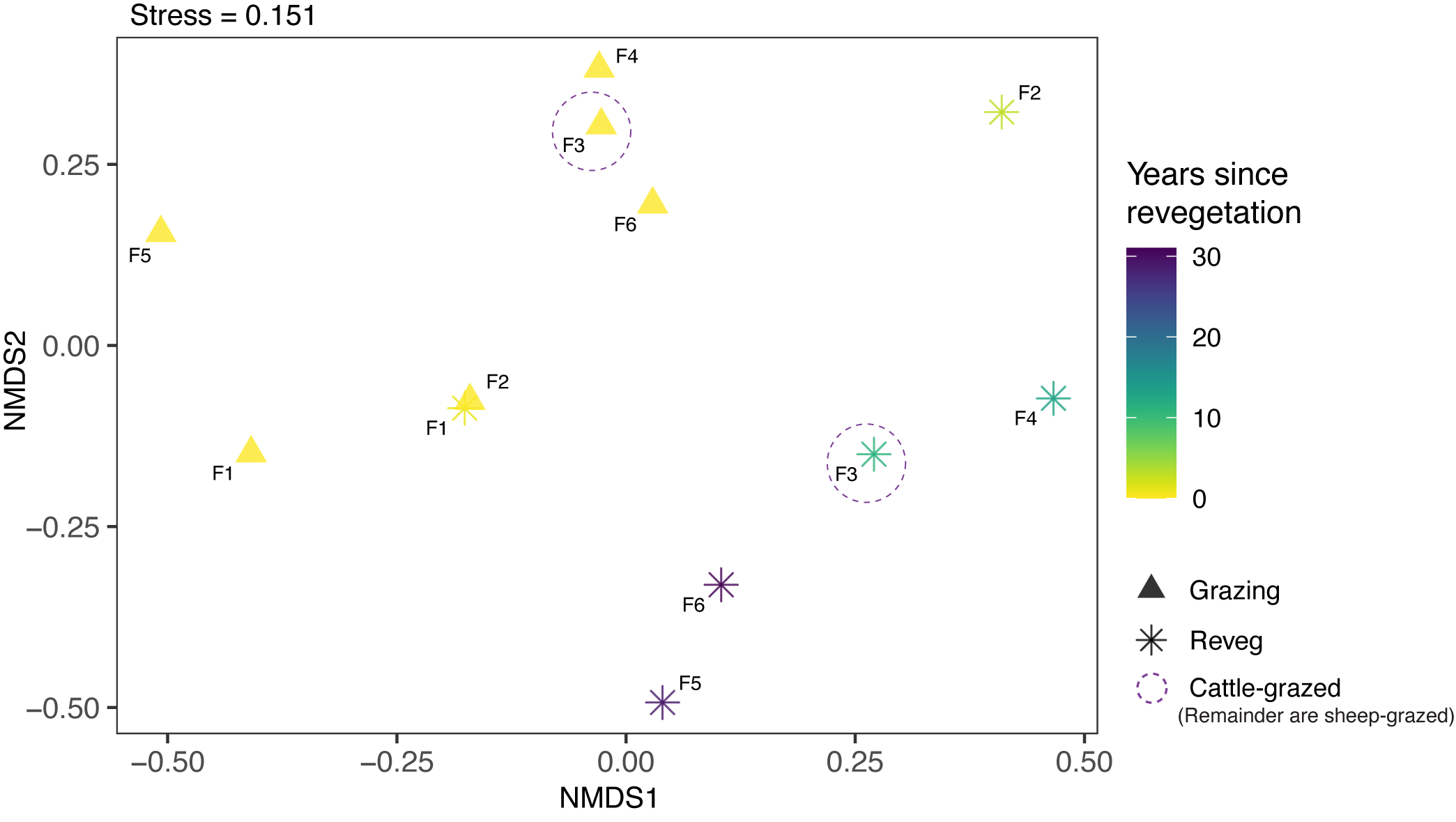

**Fig. S3. Site-level differences in plant community composition.** Non-metric multidimensional scaling (NMDS) visualising site-level differences in putative plant community composition (i.e., beta-diversity). Putative plant communities were inferred from an eDNA approach using the soil metagenomic data (see Methods). Community dissimilarities were derived from pairwise Bray-Curtis distances. Point shapes indicate land-use type: grazing (triangles) versus revegetated (stars). Points are coloured by years since revegetation. Text labels indicate the farm from which the samples derived (F1 to F6). Note, F3 represents the cattle-grazed farm, while all others are sheep-grazed.
